# Identification of Novel Common Breast Cancer Risk Variants in Latinas at the 6q25 Locus

**DOI:** 10.1101/343806

**Authors:** Joshua Hoffman, Laura Fejerman, Donglei Hu, Scott Huntsman, Min Li, Esther John, Gabriela Torres Mejia, Larry Kushi, Yuan Chun Ding, Jeffrey Weitzel, Susan L. Neuhausen, Paul Lott, COLUMBUS Consortium, Magdalena Echeverry, Luis Carvajal Carmona, Esteban Burchard, Celeste Eng, Wei Zheng, Jirong Long, Olufunmilayo Olopade, Dezheng Huo, Christopher Haiman, Elad Ziv

**Affiliations:** Department of Epidemiology and Biostatistics, Institute of Human Genetics, University of California, San Francisco, San Francisco, CA; Division of General Internal Medicine, Department of Medicine, Institute of Human Genetics, Helen Diller Family Comprehensive Cancer Center, University of California, San Francisco, San Francisco, CA; Division of General Internal Medicine, Department of Medicine, University of California, San Francisco, San Francisco, CA; Cancer Prevention Institute of California, Fremont, CA, Stanford Cancer Institute, Stanford University, Palo Alto, CA; National Institute of Public Health, Cuernavaca, Mexico; Division of Research, Kaiser Permanente, Northern California; Beckman Research Institute of City of Hope, Department of Population Sciences, City of Hope, Duarte, CA; City of Hope National Medical Center, Clinical Cancer Genetics, Duarte CA; Department of Biochemistry and Molecular Medicine, University of California, Davis, Davis, CA; University of Tolima, Ibague, Colombia; Department of Biophamaceutical Sciences, Lung Biology Center, Department of Medicine, Institute for Human Genetics, University of California, San Francisco, San Francisco, CA; Department of Epidemiology, Vanderbilt University Medical Center, TN; Department of Public Health Sciences, University of Chicago School of Medicine, Chicago, IL; Department of Medicine, Section of Oncology, University of Chicago School of Medicine, Chicago, IL; Department of Preventive Medicine, Norris Comprehensive Cancer Center, Keck School of Medicine, University of Southern California, Los Angeles, CA

**Keywords:** Genome wide association study, fine mapping, Hispanic/Latino populations

## Abstract

Background: Breast cancer is a partially heritable trait and over 180 common genetic variants have been associated with breast cancer in genome wide association studies (GWAS). We have previously performed breast cancer GWAS in Latinas and identified a strongly protective single nucleotide polymorphism (SNP) at 6q25 with the protective minor allele originating from Indigenous American ancestry. Here we report on additional GWAS and replication in Latinas.

Methods: We performed GWAS in 2385 cases and 7342 controls who were either U.S. Latinas or Mexican women. We replicated 2412 cases and 1620 controls of U.S Latina, Mexican, and Colombian women. In addition, we replicated the top novel variants in study of African American and African women and in one study of Chinese women. In each dataset we used logistic regression models to test the association between SNPs and breast cancer risk and corrected for genetic ancestry using either principal components or genetic ancestry inferred from ancestry informative markers using a model based approach.

Results: We identified 3 SNPs (p=1.9×10^-8^ - 2.8×10^-8^) at 6q25 locus not in linkage disequilibrium (LD) with variants previously reported at this locus. These SNPs were in high LD with each other, with the top SNP, rs3778609, associated with breast cancer with an odds ratio (OR) and 95% confidence interval (95% CI) of 0.75 (0.68-0.83). In a replication in women of Latin American origin, we also observed a consistent effect (OR: 0.88; 95% CI: 0.78-0.99; p=0.037). Since the minor allele was common in East Asians and African American but not European ancestry populations, we replicated in a meta-analysis of those populations and also observed a consistent effect (OR 0.94; 95% CI: 0.91 – 0.97; p=0.013).

Conclusion: The effect size of this variant is relatively large compared to other common variants associated with breast cancer and adds to evidence about the importance of the 6q25 locus for breast cancer susceptibility. Our finding also highlights the utility of performing additional searches for genetic variants for breast cancer in non-European populations.

## Introduction

Breast cancer is a partially heritable disease. Mutations in several high penetrance genes including *BRCA1*[1, 2], *BRCA2*[3] and others[4] are associated with high risk of breast cancer among carriers and explain a fraction of the heritability. Genome-wide association studies have identified over 180 common single nucleotide polymorphisms (SNPs) associated with risk of breast cancer [5-20]. The majority of these SNPs were identified in European ancestry and East Asian ancestry populations, although some unique SNPs have been identified in African American populations[21] and in Latina populations[22, 23].

Several GWAS studies have identified SNPs at 6q25 that are associated with breast cancer risk[13, 18, 20, 23-27] and mammographic density[23, 27-30]. The initial report identified a SNP in the intergenic region between *ESR1* and *CDCC170* in an East Asian population[24]. The locus was then confirmed in other populations and several additional variants were identified[11, 18, 25, 26, 31]. More recently, a fine-mapping and functional approach at this locus identified five distinct common variants associated with risk of different subtypes of breast cancer[27].

Hispanic/Latino populations are the second largest ethnic group in the U.S.[32] and yet have been understudied in genome wide association studies[33]. Latinos are a population of mixed ancestry with European, Indigenous American and African ancestral contributions[34-37]. Since there are no large studies of breast cancer in Indigenous American populations, studies in Latinos may identify novel variants associated breast cancer unique to or substantially more common in this population. We have previously used an admixture mapping approach to search for breast cancer susceptibility loci in Latinas and identified a large region at 6q25 where Indigenous American ancestry was associated with decreased risk of breast cancer[22]. Subsequently, we identified a SNP (rs140068132) that was common (minor allele frequency ~0.1) only in Latinas with Indigenous American ancestry and was associated with substantially lower risk of breast cancer, particularly estrogen receptor (ER) negative breast cancer and with lower mammographic density[23]. However, the variant we identified did not completely explain the risk associated with locus specific ancestry at 6q25 in Latinas, suggesting that other variants may account for this risk. We set out to fine-map and identify additional variants at 6q25 associated with breast cancer risk among Latinas.

## Methods

### Populations

San Francisco Bay Area Breast Cancer Study (SFBCS): The SFBCS is a population-based multiethnic case–control study of breast cancer. Cases aged 35–79 years diagnosed with invasive breast cancer from 1995 to 2002 were identified through the Greater Bay Area Cancer Registry. Controls were identified by random-digit dialing and matched on 5-year age groups. Blood collection was initiated in 1999. For this study, we focused only on cases and matched controls who self-identified as Latina or Hispanic and included 351 cases and 579 controls. Samples from this study were used as part of the initial discovery set.

GALA1: GALA1 is a family-based study (including children with asthma and their parents) of pediatric asthma in Latino Americans. We included 112 females of self-reported Mexican origin from the GALA1 study to our set of population controls. The individuals are between 11 and 42 years of age (85% are older than 20 years). Samples from this study were used as part of the initial discovery set.

Breast Cancer Family Registry (BCFR): The BCFR is an international, National Cancer Institute (NCI)-funded family study that has recruited and followed over 13,000 breast cancer families and breast cancer cases with strong likelihood of genetic contribution to disease45. The present study includes samples from the population-based Northern California site of the BCFR. Cases aged 18–64 years diagnosed from 1995 to 2007 were ascertained through the Greater Bay Area Cancer Registry. Cases with indicators of increased genetic susceptibility (diagnosis at the age of <35 years, bilateral breast cancer with the first diagnosis at the age of <50 years, a personal history of ovarian or childhood cancer and a family history of breast or ovarian cancer in first-degree relatives) were oversampled. Cases not meeting these criteria were randomly sampled46. Population controls were identified through random-digit dialing and frequency-matched on 5-year age groups to cases diagnosed from 1995 to 1998. We included 641 cases and 61 controls who self-identified as Latina or Hispanic from this study. Samples from this study were used as part of the initial discovery set.

Multiethnic Cohort (MEC): The MEC is a large prospective cohort study in California (mainly Los Angeles County) and Hawaii. The breast cancer study is a nested case–control study including women with invasive breast cancer diagnosed at the age of >45 years and controls matched on age (within 5 years) and self-identified ethnicity47. For the current study, we used data and genetic data from 546 Latina women with breast cancer and 558 matched Latina controls. We also included an additional 1,941 controls who self-identified as Hispanic/Latino from this study (935 of these controls are men) selected as part of a GWAS of type 2 diabetes[38]. Samples from this study were used as part of the initial discovery set.

Research Project on Genes Environment and Health (RPGEH): The RPGEH is a large cohort study of over 100,000 men and women of all racial/ethnic groups who are members of the Kaiser Permanente Health Plan (additional recruitment criteria?). This analysis focuses only on women who are of self-reported Latina/Hispanic ethnicity (N=3801). We included both incident and prevalent cases (total N=225) in our analyses. We identified 44 women who were also included the SFBCS. The genetic data from these participants were included as part of the RPGEH since we considered the Affymetrix Lat array as a more comprehensive array. Samples from this study were used as part of the initial discovery set.

Cancer de Mama (CAMA) Study: This study is a population-based case–control study of breast cancer conducted in Mexico City, Monterrey and Veracruz. Cases aged 35–69 years diagnosed between 2005 and 2007 were recruited from 11 hospitals (three to five in each region). Controls were recruited based on membership in the same health plan as the cases and are frequency-matched on 5-year age groups. For the current study, we used data and DNA samples from 1008 women with breast cancer and 1,063 controls. Of these 698 cases and 599 controls were genotyped with Oncoarray and included in the discovery. An additional, 310 cases and 464 controls were included as part of the replication dataset. A subset of the samples from this study were used as part of the initial discovery set and another subset were used as part of the replication.

Colombian Study of Environmental and Heritable Causes of Breast Cancer (COLUMBUS): COLUMBUS is a population-based case–control study of breast cancer conducted in four cities: Bogota, Ibague and Neiva, from the Central Colombian Andes region, and Pasto, from the Colombian South. Incident cases with invasive breast cancer aged 18–75 years have been recruited in two population registries and two large cancer hospitals. Recruitment started in 2011. Cancer-free controls were recruited through the same institutions and were matched on education, socioeconomic status and local origin using a genealogical interview. In the current study, we used data from 954 cases and 769 controls for the replication study.

Hereditary Cancer Registry of City of Hope (HCRCOH): (Southern California; PI Jeffrey Weitzel). Latina breast cancer cases are part of the HCRCOH through the Clinical Cancer Genetics Community Research Network (CCGCRN). The CCGCRN includes cancer center and community-based clinics that provide genetic counseling to individuals with a personal or family history of cancer [39]. All patients are invited to participate in the HCRCOH at the time of consultation (>90% participation). Starting in May 1998 and continuing to present, female breast cancer cases with self-reported Latino origin were seen for GCRA, enrolled in the Registry and underwent *BRCA1/2* testing after informed consent. In the current study we genotyped 1148 cases. The 347 unaffected female Latina controls were from Southern California and were invited to participate at community health fairs, flyers, and at City of Hope. These samples were used as part of the replication study.

African American Breast Cancer GWAS (AABC): The GWAS includes African American participants from 9 epidemiological studies of breast cancer, comprising a total of 3,153 cases and 2,831 controls (cases/controls: The Multiethnic Cohort Study (MEC), 734/1,003; The Los Angeles component of The Women’s Contraceptive and Reproductive Experiences (CARE) Study, 380/224; The Women’s Circle of Health Study (WCHS), 272/240; The San Francisco Bay Area Breast Cancer Study (SFBCS), 172/231; The Northern California Breast Cancer Family Registry (NC-BCFR), 440/53;The Carolina Breast Cancer Study (CBCS), 656/608; The Prostate, Lung, Colorectal, and Ovarian Cancer Screening Trial (PLCO) Cohort, 64/133; The Nashville Breast Health Study (NBHS), 310/186; and, The Wake Forest University Breast Cancer Study (WFBC), 125/153). Additional details can be found in [21, 40]. These samples were used as part of the replication study.

The ROOT consortium included six studies and a total of 1,657 cases and 2,029 controls of African ancestry: The Nigerian Breast Cancer Study (NBCS), 711/624; The Barbados National Cancer Study (BNCS), 92/229; The Racial Variability in Genotypic Determinants of Breast Cancer Risk Study (RVGBC), 145/257; The Baltimore Breast Cancer Study (BBCS), 95/102; The Chicago Cancer Prone Study (CCPS), 394/387; and The Southern Community Cohort (SCCS), 220/430. Additional details can be found in [21]. These samples were used as part of the replication study.

Shanghai Breast Cancer Study: The SBCS is a population-based, case-control study conducted in urban Shanghai. Subject recruitment in the initial phase of the SBCS (SBCS-I) was conducted between August 1996 and March 1998. The second phase (SBCS-II) of recruitment occurred between April 2002 and February 2005. Breast cancer cases were identified through the population-based Shanghai Cancer Registry and supplemented by a rapid case-ascertainment system. Controls were randomly selected using the Shanghai Resident Registry. Approximately 3500 cases and 3500 controls were recruited in the study. A subset of these including with GWAS data including 2731 cases and 2135 controls were used as part o the replication study.

### Genotyping

Genome wide association: The SFBCS, NC-BCFR, and GALA samples were all genotyped with Affymetrix 6.0 arrays at UCSF. The MEC samples were genotyped with Illumina 660 array at USC (546 Latina women with breast cancer and 558 matched Latina controls) and an additional 1941 controls were typed on an Illumina 2.5M array at the Broad Institute (Cambridge, MA). The RPGEH samples were typed on an Affymetrix LAT array at UCSF. The CAMA samples were typed on an Ilumina Oncoarray at the Quebec Genome Center. The COLUMBUS samples were typed on an Affymetrix Biobank Array. Genotyping in the AABC consortium was conducted using the IlluminaHuman1M-Duo BeadChip. Genotyping in the ROOT consortium was conducted using the Illumina HumanOmni2.5-8v1 array at Johns Hopkins University Center for Inherited Disease Research. The Shanghai Breast Cancer Study samples were typed on on an Affymetrix Genome-Wide Human SNP Array 6.0. After quality control exclusions, the final data set included 2731 cases and 2135 controls for 668 499 markers.

Replication genotyping: The CAMA samples which were not included in the GWAS and the CCGRN samples, were genotyped using Taqman probes for rs3778609. The CAMA samples included 106 ancestry informative markers from genotyped on a Sequenome platform as previously described[41]. CCGRN samples included 100 ancestry informative markers that were included as part of a sequencing project. The sequence data were aligned to Hg37 using Burrows-Wheeler Alignment and genotype calls were made using Haplotypecaller which is part of the GATK platform.

### Analysis

#### Genotyping Quality Control and Imputation

Samples with >5% missing genotypes were removed from each dataset. We dropped variants with >5% missing data from each dataset. Since excess homozygosity is more common in populations with substructure, particularly with ancestry informative markers, we did not use deviation from Hardy-Weinberg equilibrium as a criteria for excluding markers. All datasets were entered mapped to Hg19. Each dataset was then phased using SHAPEIT and imputed using the Haplotype Reference Consortium (HRC) with Minimac3 [45]. For the MEC datasets which included both 660K and 2.5M arrays, we used the overlapping SNPs (N=192,795) and imputed from those since we found that if we imputed them separately and then analyzed them together we got a large number of false positives. Each of the remaining GWAS datasets was submitted to the HRC server individually for imputation. Only variants with imputation quality scores of R2>0.5 were selected for additional analysis.

Genotype imputation for the ROOT consortium was conducted using the *IMPUTE2* software [42] with the 1000 Genomes Project phase I cosmopolitan variant set as the reference panel (October 2011 release). Genotype imputation in AABC was conducted using *IMPUTE2* software [42] to a cosmopolitan panel of all 1000 Genome Project subjects (March 2012 release). Variants with imputation score >0.3 were included in the analysis.

The Shangai Breast Cancer Study GWAS data were phased with Minimac2 and imputed with SHAPEIT using 1000 Genomes Project Phase 3. Only SNPs with an MAF ≥ 0.01 and high imputation quality (RSQR ≥ 0.5) in three GWAS in the analyses.

We used KING[43] to identify relative pairs either within the RPGEH cohort or between the RPGEH and SFBCS and/or NC-BCFR and performed the same analysis within the MEC and the CAMA study. We identified pairs of individuals with kinship coefficient >0.2 and dropped one from each of these pairs. If a relative pair included a case and control then we excluded the control. If a relative pair includes two cases or two controls we randomly dropped one of them. We dropped 127 individuals to eliminate all closely related individuals from the combined RPGEH, SFBCS and NC-BCFR.

#### Empirical Assessment of Imputation accuracy

We genotyped rs3778609, the top novel SNP, in the CAMA study in samples that also had GWAS data and checked the concordance between genotyped and imputed results. We found excellent concordance between the imputed and genotyped data with 1361/1369 (99.4%) concordance between the genotyped and imputed datasets.

#### Genetic Ancestry Inference

We implemented principal component analysis to assess genetic ancestry in each of the discovery datasets in unrelated individuals. To do so, we first LD pruned typed SNPs with r^2^ > 0.2 in PLINK. With the remaining data, we determined the principal components (PC) using EIGENSTRAT[44] within smartpca. For the replication datasets, we used ancestry informative markers and used the program ADMIXTURE[45] to calculate genetic ancestry, assuming a 3 population model with ancestry from African, European and Native American populations.

#### Association Testing

We performed single variant association testing using logistic regression models and adjusting for PC’s 1-10 in PLINK[46]. For the replication datasets we entered ancestry into the model as covariates. We also performed association testing separately for estrogen receptor (ER)-positive and ER-negative breast cancer using this approach. In each analysis we also included study as a covariate.

To calculate linkage disequilibrium (LD), we calculated R^2^ in the controls in our dataset using PLINK. We then performed conditional analyses by entering the most significant SNP in the model as a covariate in addition to PC’s 1-10.

#### Power

Based on the sample size for discovery (2396 cases and 7468 controls) we had ~80% power to detect an odds ratio of 1.25, 1.355 and 1.475 with allele frequencies of 0.4, 0.2, and 0.1 respectively.

## Results

### Individual Association Analyses

We conducted a meta-analysis across four GWAS discovery studies (Table 1) and identified 28 variants with genome-wide significant p-values at the 6q25 susceptibility region(Supplementary Table 1). No additional genome wide significant SNPs were identified. The top variants in the region included rs140068132 and rs147157845 which are in near perfect LD (r^2^=0.96) and which we have previously reported as genome-wide significant in this population[23]. Of the 28 top variants, 25 were in strong linkage disequilibrium (r^2^>0.4) with rs140068132, and 3 were in low LD (r^2^<0.2). These SNPs, rs3778609, rs7771984 and rs6914438 have a minor allele frequency of 0.19 and are in near perfect LD (r^2^=0.99). The minor allele of these SNPs are associated with lower risk of breast cancer, and the odds ratio (OR) for rs3778609 was 0.75 (95% CI: 0.68 – 0.93, p=1.9×10^-8^; Table 2). These SNPs are also independent (r^2^<0.2) of previously reported SNPs at this locus (Supplementary Table 2). Another SNP, rs851983, was associated with a near genome-wide significant level of association (MAF=0.35; OR: 1.24, 95% CI: 1.25-1.34, p= 5.6×10^-8^ Table 2). However, this SNP is in strong LD with SNPs that were previously reported (Supplementary Table 2).

**Table 1:** Discovery and Replication Samples Used

| **Discovery: Latinas** |  |  |  |
| --- | --- | --- | --- |
| Study | Genotyping Platform | Cases | Controls |
| BCFR/SFBACS | Affy 6.0 | 942 | 699 |
| RPGEH | Affy Axiom | 225 | 3574 |
| MEC | Illumina 1M, 2.5M | 520 | 2470 |
| CAMA | Illumina Oncoarray | 698 | 599 |
| **Total** |  | **2385** | **7342** |
| **Replication: Latinas** |  |  |  |
| COLUMBUS | Affy Axiom | 954 | 769 |
| CCGRN | Taqman | 1148 | 387 |
| CAMA | Taqman | 310 | 464 |
| **Total Latina replication** |  | **2412** | **1620** |
| **Replication: African American** |  |  |  |
| AABC | Illumina 1M | 3153 | 2831 |
| ROOT | Illumina 2.5M | 1657 | 2029 |
| **Total African ancestry replication** |  | **4810** | **4860** |
| **Replication: East Asian ancestry** |  |  |  |
| Shanghai Breast Cancer Study | Affymetrix 6.0 | 2731 | 2135 |

**Table 2:** Representative SNPs and association statistics from each of 4 different SNP clusters/regions that are genome wide significant.

| SNP/Risk allele | Position (BP, Hg19) | Odds Ratio (95% CI) | P value | Conditional OR* p value | Joint OR** P value |
| --- | --- | --- | --- | --- | --- |
| rs140068132-G | 6:151954834 | 0.58 (0.51-0.67) | $1.2 \times 10^{-13}$ | | 0.66, $5.1 \times 10^{-8}$ |
| rs3778609-T | 6:152133187 | 0.75 (0.68-0.83) | $1.9 \times 10^{-8}$ | 0.83, 0.0009 | 0.87, 0.007 |
| rs7771894-T | 6:152145916 | 0.76 (0.68-0.83) | $2.3 \times 10^{-8}$ | 0.84, 0.001 | |
| rs6914438-T | 6:152129588 | 0.73 (0.65-0.81) | $4.3 \times 10^{-8}$ | 0.81, 0.001 | |
| rs851983-G | 6: 152024415 | 1.24 (1.15-1.34) | $5.6 \times 10^{-8}$ | 1.17, 0.0007 | 1.15, 0.001 |

\*Conditional on rs140068132
\*\* Joint model with rs140068132, rs3778609 and rs851983

We performed conditional analyses by entering rs140068132 and other top SNPs at this locus in joint models. We found that rs3778609, rs7771984 and rs6914438 all remained nominally significant in joint models adjusting for rs140068132 (Table 2), although the adjusted odds ratios for these 3 SNPs were attenuated (OR~0.81-0.84; p<=0.001). We also found that rs851983 remained nominally significant in joint models with rs140068132 with mild attenuation. When we included 3 SNPs that best represent each of the signals from each set of associated variants (rs140068132, rs3778609 and rs851983) in the same model all of the SNPs remained nominally significant with minimal attenuation of the odds ratios (compared to models including just pairs of variants; Table 2).

### Technical Validation and Replication

We used data from the portion of the CAMA study that did not have GWAS data, The COLUMBUS study and the CCGRN to replicate the association with rs3778609. We found a consistent direction in all 3 studies and a nominally significant association in a meta-analysis of the 3 studies (n= cases; n=controls; OR=0.88, 95% CI: 0.78-0.99, p=0.037, Table 3).

**Table 3:** Replication of rs3778609 in other Latina datasets

| Study | Odds Ratio (95% CI) | P value |
| --- | --- | --- |
| COLUMBUS | 0.87(0.73 -1.04) | 0.119 |
| CCGRN | 0.89(0.70 -1.14) | 0.375 |
| CAMA (excluding GWAS) | 0.88(0.69 -1.13) | 0.314 |
| **Meta-analysis** | **0.88(0.78 -0.99)** | **0.037** |

**Table 4:** Association by Estrogen Receptor Status

| SNP | ER-positive |  | ER-negative |  | P value for ER-positive vs. ER-negative |
| --- | --- | --- | --- | --- | --- |
|  | Odds Ratio (95% CI) | P value | Odds Ratio (95% CI) | P value |  |
| rs140068132-G | 0.57 (0.46-0.72) | $1.0 \times 10^{-6}$ | 0.35 (0.23-0.54) | $2.2 \times 10^{-6}$ | 0.033 |
| rs3778609-T | 0.75 (0.65-0.86) | $6.7 \times 10^{-5}$ | 0.59 (0.46-0.75) | $2.7 \times 10^{-5}$ | 0.037 |
| rs851983-G | 1.22 (1.10-1.36) | $2.4 \times 10^{-4}$ | 1.36 (1.15-1.62) | $4.6 \times 10^{-4}$ | 0.33 |

### Association of Previously Identified SNPs at 6q25

We examined previously reported SNPs in our discovery dataset (Supplementary Table 3). Only rs851984 was significant in our study. However, nearly all of the others were directionally consistent and the 95% confidence intervals overlapped with the results from previous studies.

### Association with Estrogen receptor subtypes

We analyzed the association for each of the top SNPs separately and jointly by ER-status. As we have previously reported the minor (low risk) allele of rs140068132 is associated with a significantly lower odds ratio for ER-negative than for ER-positive breast cancer. We also found a significantly stronger effect size for ER-negative breast cancer for rs3778609. The effect size for rs851983 is also greater for ER-negative breast cancer; however, there is no significant difference between ER-negative and ER-positive breast cancer for this SNP.

### Replication in non-Latinas

The cluster of new SNPs we identified in this locus represented by rs37786109 are common in East Asians (minor allele frequency 0.27) and African (minor allele frequency 0.3) ancestry populations (Supplementary table 4), but not in European populations (minor allele frequency 0.019 in 1000 Genomes). Therefore, we evaluated the association with this SNP in several additional populations in patients of these ancestries including the African American Breast Cancer study, the ROOT study, and the Shanghai Breast Cancer Study. We found a consistent effect for of these studies with a significant effect in a meta-analysis of rs3778609, rs7771894 and rs6914438 (Supplementary table 4). We also examined rs3778610 which is in strong LD with these SNPs and had near genome-wide significant associations in our discovery sample (Supplementary Table 1). The strongest association in non-Latina populations was with rs3778610 which was particularly stronger in African ancestry populations (Supplementary table 4).

## Discussion

We have previously reported on a SNP at 6q25 associated with a minor allele that is unique to Indigenous American populations and associated with decreased risk of breast cancer [23]. Here, we investigate this locus in greater depth in an expanded sample size of Latina breast cancer cases and controls, the largest sample size of this population analyzed for breast cancer risk to date. We have identified several SNPs that are genome-wide significant and associated with breast cancer at this locus independently of other SNPs at this region previously reported by us and those previously reported by other groups at this locus. These SNPs are located in the region of ~152.13 – 152.14 MB (Hg19). Replication in African American and Asian samples supports the association with this SNP. In addition, we have also shown that these novel variants at this locus have significantly stronger effect sizes on ER-negative breast cancer. Prior studies have also demonstrated a stronger effect size with ER-negative breast cancer for most variants at the 6q25 locus, consistent with our data [18, 27].

Prior studies in other populations have reported a series of independent SNPs affecting breast cancer risk [11, 18, 24-27, 31]. The variants previously reported in other populations are not significant in our studies but most are directionally consistent. A combined fine-mapping and functional study of SNPs mapped in other populations at this locus found that they affect expression of *ESR1*, *RMND1* and *CCDC170[27]*. Since the new variants we report are common only in non-European ancestry populations, there is limited data to explore the potential effects of these variants on gene expression.

Our study is limited by sample size. Therefore, it is possible that we have missed other variants at this locus. In fact, several previously reported variants have odds ratios with point estimates that are close to the previously reported results, but have 95% confidence intervals that include 1, as would be expected with insufficient power. The effect size we observed in the replication dataset is substantially lower than in the discovery dataset, likely due to winner’s curse. However, even if we take the replication odds ratio (0.88) as the closest to the true effect size of these SNPs, this is still a relatively large effect for a common variant. It is likely that there are other variants that have not yet been identified in European GWAS due to low allele frequency and that could be identified in Latinas where they are more common. Larger studies of Latina women are needed to identify these variants.

## Conclusion

Our study demonstrates additional unique associations with variants at 6q25 and breast cancer risk. This further highlights the important contribution of this locus to breast cancer susceptibility, particularly ER-negative breast cancer susceptibility. Additional fine- mapping and functional studies are needed to elucidate all of the causal variants in our population. However, the variants we identified in this study can be useful to add to the increasing pool of common variants coming from GWAS and will be particularly useful to risk stratify women of Latin American ancestry for breast cancer risk.

## List of abbreviations

GWAS: Genome wide association study
SNP: Single nucleotide polymorphism
OR: odds ratio
CI: confidence interval
PCA: Principal components analysis
PC: Principal component
ER: Estrogen receptor

## Declarations

Ethics approval and consent to participate: The following Institutional Review Boards have approved the collection of different datasets which have contributed to this study: University of California, San Francisco, the Cancer Prevention Institute of California, University of Southern California, Kaiser Permanente Division of Research Northern California, University of Chicago, Vanderbilt University, National Institute of Public Health, Mexico

### Consent for publication

Not applicable

### Availability of data and material

The following GWAS datasets have been submitted to dbGAP: SFBACS (phs000912.v1.p1), MEC Latina (phs000517.v2.p1), AABC (phs000851.v1.p1), ROOT (phs000383.v1.p1), Shanghai Breast Cancer Study (phs000799.v1.p1). The NC-BCFR was determined by the IRB not to be eligible for submission to dbGAP based on the consent. Summary statistics are available for the 6q25 locus from the NC-BCFR, the Kaiser RPGEH and the COLUMBUS study are available from the corresponding author by request.

### Competing interests

The authors declare no competing interests.

### Funding

This work was funded in part by Grants from the National Cancer Institute (K24CA169004, R01CA120120 to E Ziv and R01CA184545 to E Ziv and S Neuhausen). J Hoffman was supported by R25CA112355. The San Francisco Bay Area Breast Cancer Study was supported by the National Cancer Institute (CA63446 and CA77305), the US Department of Defense (DAMD17-96-1-6071) and the California Breast Cancer Research Program (7PB0068). The Northern California Breast Cancer Family Registry was supported by grant UM1 CA164920 from the National Cancer Institute. The Multiethnic Cohort Study was supported by the National Institutes of Health grants R01 CA63464 and R37CA54281, R01 CA132839, 5UM1CA164973. The CAMA Study was funded by Consejo Nacional de Ciencia y Tecnología (SALUD-2002-C01-7462). The COLUMBUS study receives support from GSK Oncology (Ethnic Research Initiative), University of Tolima, University of California Davis and The V Foundation. L.C.-C. is a V Foundation V Scholar. The Shanghai study was supported by R01CA148667 and UM1 CA182910. The ROOT study was supported by the American Cancer Society (MRSG-13-063-01-TBG and CRP-10-119-01-CCE), National Cancer Institute (CA142996, CA161032), Susan G. Komen for the Cure, and Breast Cancer Research Foundation. The GALA study was supported by the Sandler Family Foundation, the American Asthma Foundation, the RWJF Amos Medical Faculty Development Program, Harry Wm. and Diana V. Hind Distinguished Professor in Pharmaceutical Sciences II, National Institutes of Health 1R01HL117004, R01Hl128439, R01HL135156, 1X01HL134589, National Institute of Health and Environmental Health SciencesR01ES015794, R21ES24844, the National Institute on Minority Health and Health Disparities 1P60MD006902, RL5GM118984, 1R01MD010443 and the Tobacco-Related Disease Research Program under Award Number 24RT-0025.

### Authors’ contributions

JH, LF and EZ conceived of the study. JH, LF, DH, SH conducted GWAS analyses in discovery populations. ML and YCD conducted replication genotyping. CH contributed samples and data, contributed to analyses and made critical comments to the manuscript. EJ, GTM, LK, SN, JW, OO, DH, JL, WZ, PL ME LCC, EB, CE contributed samples and data and made revisions to the manuscript.

